# Slow Oscillations Modulate Functional Brain Changes Supporting Working Memory

**DOI:** 10.1101/2024.05.14.594161

**Authors:** Jing Zhang, Pin-Chun Chen, Sara C. Mednick, Arielle Tambini

## Abstract

Working memory (WM), the temporary mental storage and manipulation of information, is a skill that can improve with training. Sleep, and specifically slow oscillations (SOs), has been linked with WM improvement, yet it is unknown how processing during SOs modulates WM function across sleep. The current study examines how WM-related neural processing changes with sleep, and how these changes are related to activity during SOs. To do so, participants performed a WM task during fMRI before and after sleep, and the first 2.5 hrs of sleep was monitored by simultaneous EEG-fMRI. Reliable overnight changes in WM-related activity patterns were found, with reduced recruitment of the dorsal precuneus after compared to before sleep. Moreover, greater neural activation during SOs was associated with reduced overnight recruitment during WM across multiple levels of analysis. Our findings highlight the important role of SOs, elucidating how SOs may support changes in WM function across sleep.

## Introduction

Working memory (WM) is a critical aspect of cognition that facilitates temporary storage and manipulation of information to achieve a goal. Beyond temporarily helping you remember a phone number until you can write it down, WM also supports a wide range of cognitive functions that are fundamental to many everyday tasks, such as decision-making, problem-solving, medication adherence, and language comprehension^1–4^. Given its functional importance and its vulnerability to age-related decline^5^, cognitive training to improve WM function has become a focus of research. The hope is that by improving WM, executive functions will also improve, which could mitigate cognitive aging. Several studies have demonstrated that a period of sleep, compared to wake, between WM sessions supports WM improvement^6–9^. And yet, the neural mechanisms underlying sleep-dependent WM improvement are unknown. The current study investigates this question.

Studies have demonstrated that WM capacity can be enhanced through training and is reflected in plasticity in WM-related brain activity patterns^10–12^. Specifically, Jaeggi and colleagues trained participants on a WM task for multiple days, and found a positive correlation between the extent of WM improvement and number of training sessions, as well as transfer of these benefits to a measurement of fluid intelligence^11^. This improvement in WM may be mediated by changes in brain activity induced by WM training, including increased activation in the prefrontal^13^ and parietal regions^14^, and decreased functional connectivity within the striatal-prefrontal network^15^. Furthermore, sleep plays a role in practice-induced changes in WM circuits, with greater behavioral improvement after a night of sleep^7,16^ or a nap^8,9,17^ compared to wake. On the other hand, sleep deprivation or restriction leads to negative impact on WM function, such as performance on digit span^18^ and N-back tasks^19^. These effects are likely driven, in part, by an influence of sleep on the function of frontal and parietal cortices^20^, though the neural processing during sleep that supports changes in WM function across sleep have not been directly examined.

Several studies have reported a specific role of slow wave sleep (SWS) in cognitive processing via its role in large scale, network-level plasticity^21–23^. For example, the duration of SWS has been related to improvements in digit span backward performance across sleep^24^. Slow wave activity (SWA), the slow, high-amplitude waves (<4Hz) that are observed in the electroencephalogram (EEG) during SWS, reflects synchronized population-level neuronal activity^25,26^ and has been associated with WM improvement. Specifically, after three weeks of WM training, increases in prefrontal SWA correlated with WM performance improvement^27^. Furthermore, SWA after training predicted WM gains across a period of sleep in both young^28^ and older adults^17^. Importantly, SWA may play a potentially causal role in WM improvement, as participants with greater SWA induced by an acoustic slow-wave stimulation during nighttime sleep showed better WM improvement, compared to individuals who did not show enhanced SWA^29^. In addition, slow oscillations (SOs, <1Hz), which is a type of SWA with a frequency of less than 1Hz associated with the synchronization of neural activity, has been associated with WM improvement. SOs that are coupled with changes in heart rate positively correlate with WM improvement overnight^30^. Taken together, these studies support the role of sleep, specifically SOs and SWA during SWS, in WM improvement.

This study aims to investigate neural mechanisms of sleep-related WM improvement by measuring changes in online WM brain activity over a night of sleep, and relating these changes to neural processing during sleep, specifically SO windows^31^. We tested the operational span (OSPAN) task, which measures an individual’s ability to maintain a set of items in their WM while performing a secondary math task. Prior work has shown that the OSPAN task is sensitive to changes in WM function across sleep^17,32^, which is why we chose this task as opposed to other standard WM tasks (e.g. delayed match to sample). Specifically, in the evening, participants performed the OSPAN task during fMRI and then fell asleep while their brain activity was recorded using simultaneous EEG-fMRI (Figure 1a). In the morning, participants were retested on the OSPAN task during fMRI. To increase the robustness of our data, each participant performed two sessions, with a 2-week washout period between sessions. We explicitly examine associations between overnight changes in online WM function/activity and SO activity. To do so, our analysis took two approaches, a data-driven analysis that examined how WM-related brain activity changed across the night; and *a priori* network-driven analysis that focused on the Dorsal and Ventral Attention Networks^33,34^.

**Figure 1.**
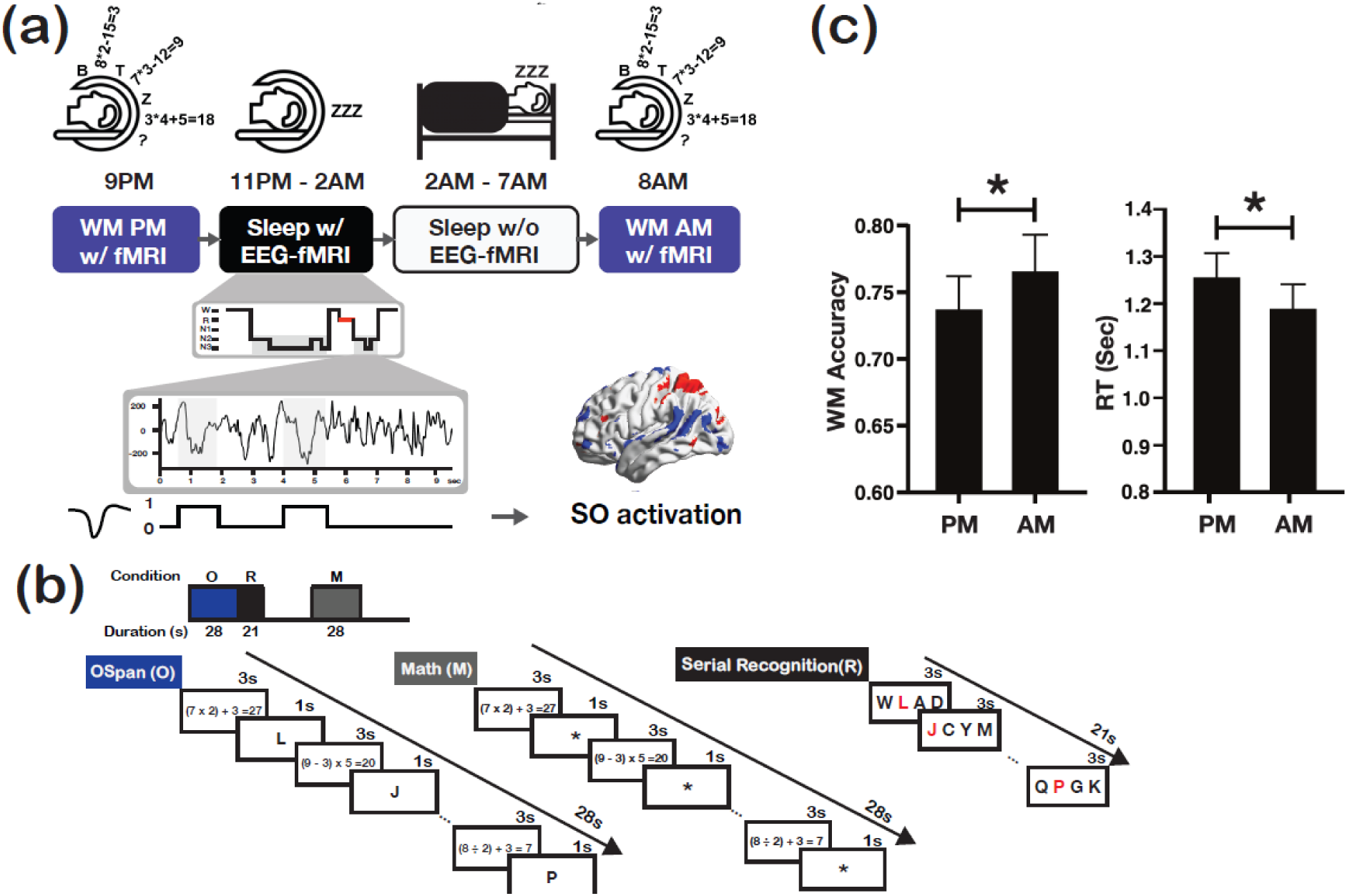
(a) Timeline of the study visit. (b) An illustration of the WM tasked used in the scanner. (c) WM accuracy for PM (M=0.74± SE 0.03) and AM (M=0.77± SE 0.03) test (left), WM reaction time (RT) for PM (M=1.26± SE 0.05 s) and AM (M=1.19± SE 0.05 s) test (right). Error bars indicate standard error (SE).

WM engages a distributed network of brain regions, including portions of the prefrontal cortex, parietal cortex, and the precuneus^35–40^. Studies have shown the involvement of two distinct attention networks supporting WM: the dorsal attention network, comprised of the intraparietal sulcus (IPS) and the superior frontal gyrus (SFG), and the ventral attention (or salience) network, including the temporoparietal junction (TPJ), inferior frontal cortex, the anterior insula, and the dorsal anterior cingulate cortex^41–43^. The dorsal attention network is associated with top-down task-focused attention and WM load^33^. The ventral attention (or salience) network is thought to govern bottom-up attention, typically engaged by stimuli that are novel or salient, as well as switching between internal and external attention^44–47^. Here, in addition to using a data-driven approach to isolate overnight changes in WM-related activation and examine links with SO activity, we also investigated links between overnight WM function and SO-related brain activity in these established networks.

To our knowledge, this is the first study to examine how functional changes time-locked to SOs contribute to neural processing during WM (e.g., overnight changes in WM-related brain activity) using simultaneous EEG-fMRI. By addressing these questions, we hope to gain a deeper understanding of the neural mechanisms underlying WM improvement and the role of sleep in shaping WM function.

## Methods

### Participants

The study complied with the ethical standards of the relevant national and institutional committees on human experimentation and with the Helsinki Declaration of 1975, as revised in 2008, and was approved by the University of California, Irvine, IRB committee (HS# 2020-5912). Sixty healthy females (18-40y) signed informed consents. Sixteen participants withdrew from the study before the first experimental visit. Four participants withdrew during the first experimental visit, resulting in 40 participants completing the first visit. Six participants were excluded after the first experimental visit due to an inability to sleep in the scanner. In total, 34 participants (M_age_ = 25.91 ± 5.29 years, all right-handed) completed the entire study protocol consisting of two overnight visits. Participants had no current or past history of major psychological or medical conditions or sleep disorders, and additionally were not taking any medications (e.g., hypnotics) that would affect cognitive function or sleep. All participants included in the study reported consistent sleep habits in advance of the study, with sleep times before 1 am and wake times before 10 am. Participants completed the Morningness-Eveningness Questionnaire (Horne & Östberg, 1976) and Epworth Sleepiness Scale (ESS) as screening assessments, with an acceptable range between 6-21 for Morningness-Eveningness Questionnaire and <15 for ESS.

### Procedure

All participants completed an orientation session to give consent and practice the OSPAN task, which occurred on a day prior to the experimental visits. During the practice session, participants practiced the OSPAN task by viewing it on a computer monitor and responding as they would with the response keypads in the MRI scanner. After task practice, participants spent 15 minutes in a mock scanner to get familiarized with sleeping in the scanner. On each experimental visit, the participant arrived at 8 PM to undergo a T1-weighted structural scan. Participants then performed the OSPAN task, followed by a paired associate learning task (unrelated to the OSPAN task and beyond the scope of the current paper), while fMRI was performed. Participants were then removed from the MRI scanner prepared for bedtime, after which the EEG cap was placed. At around 11 pm, participants were placed in the MRI scanner and had a 2.5h sleep opportunity with continuous EEG-fMRI recording (Table 1). Participants were provided with earplugs (∼33 dB sound attenuation), comfortably bedded on a viscoelastic mattress, and covered with a light blanket. The lights were turned off and subjects were equipped with an alarm bell to alert the experimenter and terminate the scan at any time. After awakening, subjects spent the remaining night in the adjacent sleep laboratory, during which their sleep was monitored using OURA ring (Table 2). Participants were woken up around 7:30 AM the next morning and returned to the scanner around 8 AM to perform the OSPAN task. The study visit timeline is shown on Figure 1a.

**Table 1.**
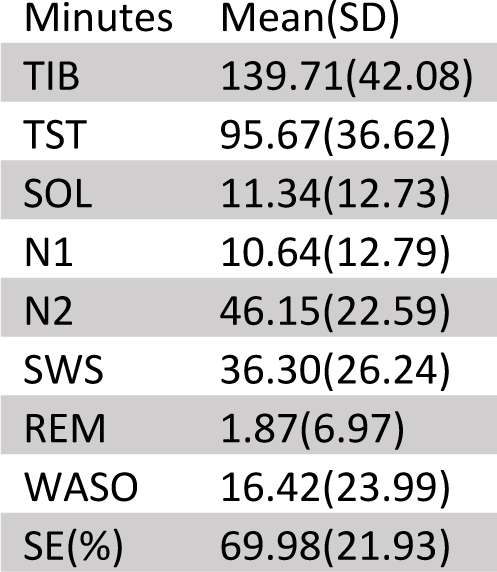
Sleep architecture during EEG-fMRI recording. Means ± SD. TIB, time in bed; TST, total sleep time; SOL, sleep onset latency; N1, stage 1 sleep; N2, stage 2 sleep; N3, stage 3 sleep/slow-wave sleep; REM, nonrapid eye movement sleep; WASO, wake after sleep onset; SE, sleep efficiency.

| Minutes | Mean(SD) |
| --- | --- |
| TIB | 139.71(42.08) |
| TST | 95.67(36.62) |
| SOL | 11.34(12.73) |
| N1 | 10.64(12.79) |
| N2 | 46.15(22.59) |
| SWS | 36.30(26.24) |
| REM | 1.87(6.97) |
| WASO | 16.42(23.99) |
| SE(%) | 69.98(21.93) |

**Table 2.** Sleep architecture total (in scanner + out of the scanner, measured by OURA)

| Minutes | Mean(SD) |
| --- | --- |
| TIB | 493.24(73.78) |
| TST | 366.06(108.39) |
| Light | 197.28(58.12) |
| Deep | 140.01(46.07) |
| REM | 74.33(132.95) |
| WASO | 49.46(32.01) |

### Task and stimuli

The task was programmed using MATLAB UI Figure; stimuli were displayed using a BOLDScreen32 LCD monitor (1920×1080 pixels with a 120Hz framerate). Responses and response times (RT) were recorded using MR compatible response keypads. Participants performed the OSPAN task in a similar fashion to Faraco et al. (2011)^48^. Each run consisted of fixed alternating conditions of OSPAN (O), Math (M), and Arrows, each lasting 28s. Task illustrations are shown in Figure 1b. During the OSPAN condition, participants judged if math equations were correct or incorrect (equations were shown for 3 s). Between equations, they were presented with a letter to remember in serial order (letters shown for 1 s). A total of seven letters were shown in each 28s OSPAN block. The OSPAN blocks were always followed by 21 s Serial Recognition (R) epochs, in which participants identified the letter shown in each serial position (1-7), by choosing between four letters across seven prompts. During the Math condition, participants also viewed math equations and judged if each was correct, but an asterisk was shown in between the equations instead of a letter (shown for the same duration as the letter). Thus, the OSPAN and Math conditions only differed based on the need to attend to and maintain serially presented letters. The Arrows condition required participants to indicate if an arrow was pointing to left or right. Each fMRI run contained a total of 17 conditions, such that the conditions: OSPAN + Serial Recognition + Arrows + Math + Arrows were repeated three times, ending with one block of the OSPAN followed by Serial Recognition. Participants performed two runs back-to-back during each session.

### WM Task Behavioral Analysis

To assess changes in WM performance across sleep, accuracy and response time (RT) during the OSPAN recognition conditions was measured. We calculated the proportion of accurate responses in each OSPAN condition as: number of correct letters recognized in the correct order divided by total number of letters in each OSPAN condition (7); therefore, the maximum score was one. WM performance was evaluated by averaging accuracy across both runs and both visits for each participant. A repeated measures ANOVA with within-subjects factors of time of day (AM/PM) and visit (1 or 2) was performed to test whether overnight changes in accuracy were reliable and varied across visits. A follow-up paired t-test was conducted between AM and PM sessions to assess sleep-related changes in WM performance (Figure 1c). To further probe changes in WM across sleep, we calculated RT during the OSPAN recognition conditions for the correct trials. RT was measured as the timing between the stimulus presentation time to response. Similar to accuracy, a repeated measures ANOVA was conducted on RT, using within-subjects factors of time of day and visit. Averaged RT was compared paired t-test between AM and PM sessions using a paired t-test.

### MRI Acquisition

Participants were scanned at the Facility for Imaging and Brain Research at the University of California, Irvine on a 3 T Siemens Prisma MRI scanner (Siemens Medical Solutions), with a 32-channel head coil. High-resolution T1-weighted anatomical images were obtained using a standard MPRAGE sequence (TR = 2300 ms, TE = 2 ms, flip angle = 8°, sagittal slices, 0.8 × 0.8 × 0.8 mm voxel size, field of view = 256 × 256 mm). Blood Oxygen Level Dependent (BOLD) functional scans (T2*-weighted) were acquired using an echo planar imaging (EPI) sequence (TR = 2240 ms, TE = 30 ms, flip angle = 90°, FOV = 216 × 216 mm, 40 transversal slices, slice thickness = 3 mm, dist factor = 10%, 3.4 × 3.4 × 3 mm voxel size, continuous bottom-up slice acquisition order). These parameters were chosen to ensure that residual gradient artifacts (in the slice repetition rate of ∼ 17 Hz) did not compromise the sleep spindle frequency. Each run of sleep scan contains maximum of 4096 volumes (approx. 2.5hrs) restricted by scanner limit. Participants were free to terminate the scan at any time and they may choose to continue sleeping in the scanner if the scan ended prematurely (see suppl. Table 1 for # of TRs obtained for each participant). Each run of the OSPAN task consisted of 216 volumes (approx. 8 minutes) and 4 dummy samples were discarded during the initial acquisition to account for equilibrium effects.

### fMRI Preprocessing

EPI volumes were slice-time corrected, co-registered, realigned, and transformed to Montreal Neurological Institute (MNI) space using fMRIPrep 1.4.0, with several confounding time-series calculated (see supplemental methods for more details; https://github.com/nipreps/fmriprep). All fMRI analyses were performed in MNI space. The fMRI data were then additionally preprocessed (spatially smoothed using a 5 mm full-width at half-maximum Gaussian smoothing kernel) and analyzed using SPM12 (Wellcome Trust Centre for Neuroimaging, London, UK, http://www.fil.ion.ucl.ac.uk/) and custom code in MATLAB (MathWorks, Natick, Massachusetts, USA).

### EEG Acquisition

EEG was acquired with an MR-compatible 12-channel EEG cap (BrainCap MR, Easy-Cap, Munich, Germany) according to the 10–20 system (F3, F4, C3, C4, O1, O2, M1, M2). Skin resistance was kept below 5 kΩ (plus 5 kΩ safety resistors) using Abralyt HiCl electrode paste (Easycap) to ensure stable EEG recordings during the 2.5 hr sleep opportunity. Bipolar recordings of electrooculogram (outer canthi), electromyogram (chin), and electrocardiogram (on the backbone, ∼ 25 cm below and above the heart) were acquired via MR-compatible Ag–AgCl cup electrodes with 10 kΩ safety resistors (EasyCap). FCz was used as the reference electrode and Cz was used as the ground electrode. The data were recorded using BrainAmp MR plus DC and bipolar BrainAmp ExG MR amplifiers and the BrainVision Recorder (BrainProducts, Munich, Germany) with a resolution of 0.5 μV/bit at 5 kHz and filtered between 0.016 and 250 Hz. To ensure precise timing between EEG and fMRI data, we used the SyncBox (Brain Products, Munich, Germany), which synchronizes the clock of the BrainAmp MR amplifier with the clock driving the MRI scanner’s gradient switching system.

### EEG Preprocessing

Online MRI gradient and ballistocardiographic artifact correction (BrainVision RecView V.1.2; BrainProducts) allowed online sleep monitoring. Removal of these artifacts for subsequent analyses was performed offline by adaptive template subtraction^49,50^ using windows of 100 volumes and 50 pulses, respectively (BrainVision Analyzer V. 2.0; BrainProducts). EEG data were re-referenced to contralateral mastoids (M1 and M2) and sleep stages were visually scored in 30-second epochs by one of the authors, according to standard American Academy of Sleep Medicine (AASM) criteria. To ensure the NREM sleep analyzed further were continuous stable NREM without transitions, EEG epochs with movements or arousal were scored as ‘motion artifacts’ instead of N2 or N3.

### Slow Oscillations (SO) Detection

Individual SO events were detected in the EEG data and then used as predictors in event-related fMRI analyses to assess BOLD activation during individual SO events (similar to^31^). SO troughs were detected for each channel automatically on MATLAB using the algorithm introduced by^51^ (also used in^30,52^). EEG data were first band pass filtered within the SO frequency range (0.7-1.4Hz) using a Hamming windowed FIR filter. The Hilbert transform was then applied to find phase and amplitude of the SO. Troughs (down-states) were detected when the phase difference > 6. The up-states before and after each trough were then detected within 1.5 seconds window before and after the troughs. SO events were defined as the pre-trough up-states of SOs (event onset) through the post-trough up-states of SOs (defined as event offset)^53–55^ during stable 30s epochs of NREM sleep (see definition in EEG Processing above). We detected on average 6.84 ± 7.98 SOs per minute across all of NREM sleep. A table of SO count and density by subject is included in supplementary materials.

### fMRI Analysis

General Linear Models (GLMs) were used to assess fMRI activation and functional connectivity. First-level analyses (model specification and estimation) were performed using SPM12. Scans were corrected for low-frequency trends in the GLM (high-pass filter with cut-off period = 111.111 seconds). Head motion was measured using framewise displacement (output from fMRIPrep). Excess variance in the BOLD signal associated with motion was accounted for by including nuisance regressors for each time point surrounding motion events (one TR before through to two TRs after FD > 0.5mm)^56^. Runs with excessive motion were not included in further analysis (% of excluded volumes greater than two standard deviations from the mean across all OSPAN task runs), which is consistent with the criterion used for sleep scans. One participant was excluded due to motion, and 3 participants had one session removed (either pm or am). Nuisance regressors were included as covariates in all GLMs: high motion volumes (defined above), six motion parameters (translation x y z; rotation x y z), the 1st temporal derivative of the six motion parameters, and the top eight anatomical CompCor (WM, CSF) components^57^ (output from fMRIPrep).

### WM-related BOLD Activation

Regressors were created to model BOLD activation during OSPAN, Math and Arrows blocks (in addition to the nuisance regressors describe above, fMRI Analysis). All 3 blocks had the same duration (28s), and onset times were calculated in 28s increments as the blocks alternated in their presentation. To assess WM-related processing, we contrasted activation during OSPAN blocks with Math blocks in each session, generating separate contrast maps (beta estimates), separately for each AM and PM session. The OSPAN vs Math contrast yielded regions associated with the encoding and maintenance of letters during the OSPAN, but not the Math, blocks, as well as additional cognitive control required during OSPAN vs. Math blocks.

To assess WM-related activation regardless of the time of day or sleep (Figure 2a), the OSPAN vs Math contrast beta maps were averaged across all four sessions (PM and AM sessions X two experimental visits) within each participant. These average beta maps were then submitted to group-level analyses (described below).

**Figure 2.**
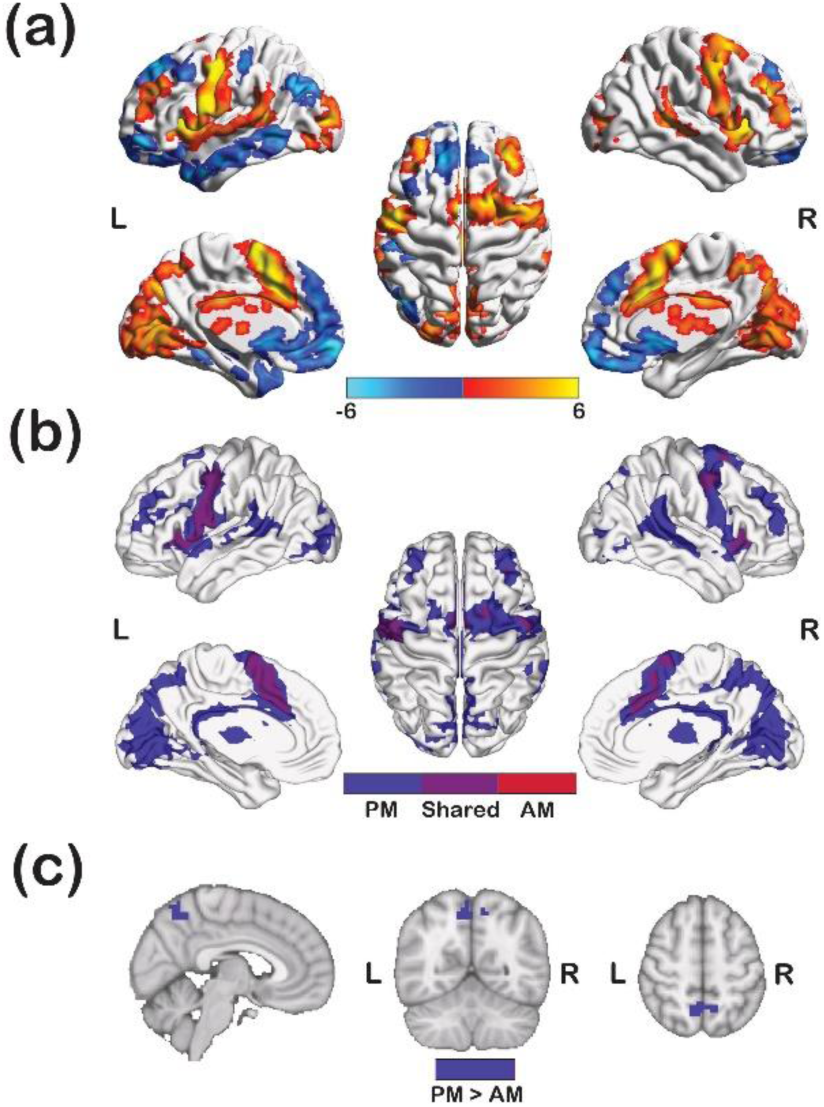
(a) Whole brain analysis on OSPAN vs Math contrast across all 4 sessions. Warm color indicates increased activation during OSPAN compared to Math and cold color indicates decreased activation. (b) WM-related activation during PM (blue), AM (red) and shared (purple). (c) The cluster in the dorsal precuneus region (MNI: 29.5 22.2, 40.4) with a significant decrease in activation overnight.

To investigate overnight changes in WM-related processing, we used two complementary approaches. First, we asked whether the pattern of overnight activation change between AM and PM sessions was reliable across the two visits. To do so, we calculated the correlation of the pattern of AM-PM changes across gray matter voxels between the two visits in each participant. Then, to test whether the similarity (i.e. reliability) of AM-PM changes across visits was robust at the group-level (i.e. consistently greater than zero), we conducted a one-sample t-test on the Fisher Z-transformed correlation (r) values across participants. Second, we localized reliable changes in WM-related activation between AM and PM sessions using a standard group-level analysis approach. Specifically, to characterize pre vs. post-sleep WM-related activation (Figure 2b), pre-sleep (PM) and post-sleep (AM) session contrast beta maps were averaged across the two visits within each participant, and sleep-dependent OSPAN changes (AM vs. PM) were calculated as the AM average – PM average within each participant. These beta maps of the AM – PM WM-related activation change were then submitted to group-level analyses (described in the paragraph below).

The contrast beta maps of WM-related activation (average across all sessions, or pre vs. post sleep changes in a separate analysis) were submitted to group-level analyses using one sample t-tests. Family-wise error (FWE) correction was performed using Randomise (https://fsl.fmrib.ox.ac.uk/fsl/fslwiki/Randomise/) with TFCE (threshold-free-cluster enhancement) and 50,000 permutations. Results are shown at the *p* < 0.05 level (FWE-corrected). The cerebellum was excluded from all analyses. We first defined data-driven WM areas as regions showing significantly greater OSPAN vs. Math activation from the group-level analysis, which showed WM-related activation regardless of the time of day (Figure 2a positive clusters). We next asked whether regions engaged during WM (OPSAN > Math) showed overnight changes in activation by testing AM vs. PM changes within the data-driven WM mask (FWE-corrected). We then further probed regions showing significant AM vs. PM WM-related activation changes (Figure 2c) by assessing their functional connectivity during the OSPAN task and during SOs (using isolated regions as seeds in Psychophysiological Interaction (PPI) analyses).

### Sleep BOLD Activation

An event-related design was used to model brain activity associated with SO events vs. periods of non-SO NREM (N2+N3 sleep). Within each run of sleep scans, scans were corrected for low-frequency trends (’spm_filter’ function; high-pass filter with cut-off period = 111.111 seconds). Next, to limit our analysis to stable NREM (N2+N3 sleep), artifact-free (as defined from the 30s segments of EEG) epochs of NREM sleep with a duration > 100 TRs (224 seconds) were extracted from the BOLD fMRI data and concatenated together (after low-frequency trends and the mean BOLD signal was removed). GLMs were performed to estimate activation associated with SO events using the concatenated, artifact-free NREM data. SO event onsets were defined as the pre-trough up states of the SOs, and event offsets were defined as the post-trough up states of the SOs (described above in SO Detection). A regressor was created to model activation related to SO events, based on these onsets and offsets. Nuisance regressors described above were included in these GLMs. The SO contrast (versus implicit baseline of non-SO NREM sleep) (Figure 4) yielded regions showing differential activation during SOs relative to overall NREM processes.

### Psychophysiological Interactions (PPI) analyses

To further probe regions showing overnight changes in the OSPAN task, a PPI approach was used to assess connectivity of these regions during WM (OSPAN vs. Math blocks). A PPI GLM (implemented in SPM12) was built using the seed region(s), chosen as all ROI(s) showing significant overnight OSPAN vs. Math activation changes (shown in Figure 2c). The PPI GLMs included one regressor representing the task contrast (OSPAN vs. Math), one regressor representing the seed ROI time series, one regressor representing the PPI term (interaction between task and seed time series), with the same nuisance covariates used as in the GLMs described above. Group-level one sample t-tests were performed on beta coefficient maps corresponding to the PPI term. As described above, FWE-correction was performed using Randomise with TFCE (Figure 3a). To measure functional connectivity of these seed region(s) during NREM sleep (SO events), the same PPI approach was applied to the NREM sleep data to assess differential connectivity during SO events. This was done by creating one regressor representing the SO events (SO vs implicit baseline of remaining NREM), one regressor representing the seed region, and one regressor representing the PPI term (Figure 3b). The cerebellum was excluded from all analyses.

**Figure 3.**
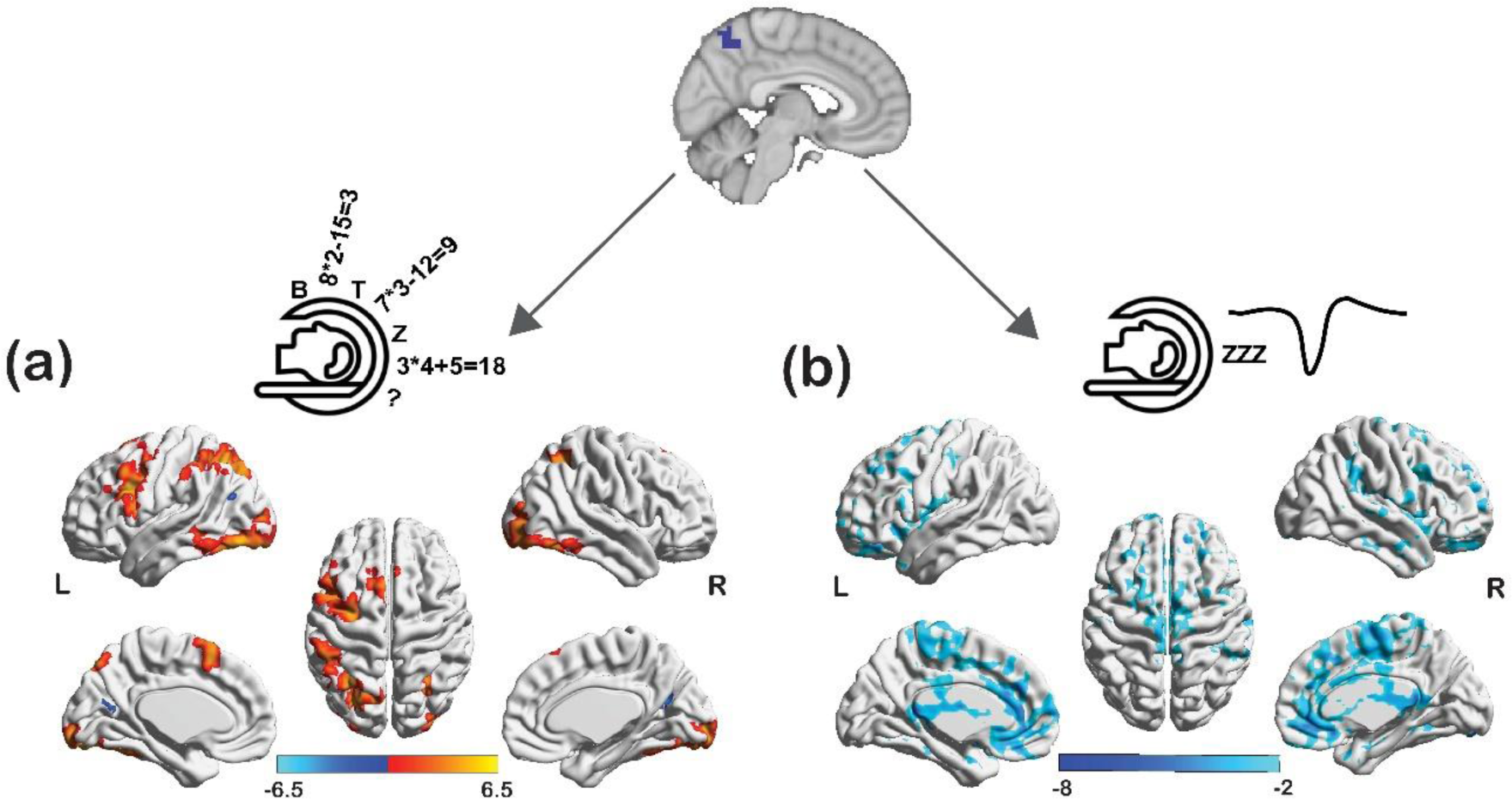
(a) Functional connectivity between the target ROI and other WM regions during OSPAN vs Math, warm color indicates increased connectivity and cold color indicates reduced connectivity. (b) Functional connectivity between the target ROI and WM regions during SOs.

**Figure 4.**
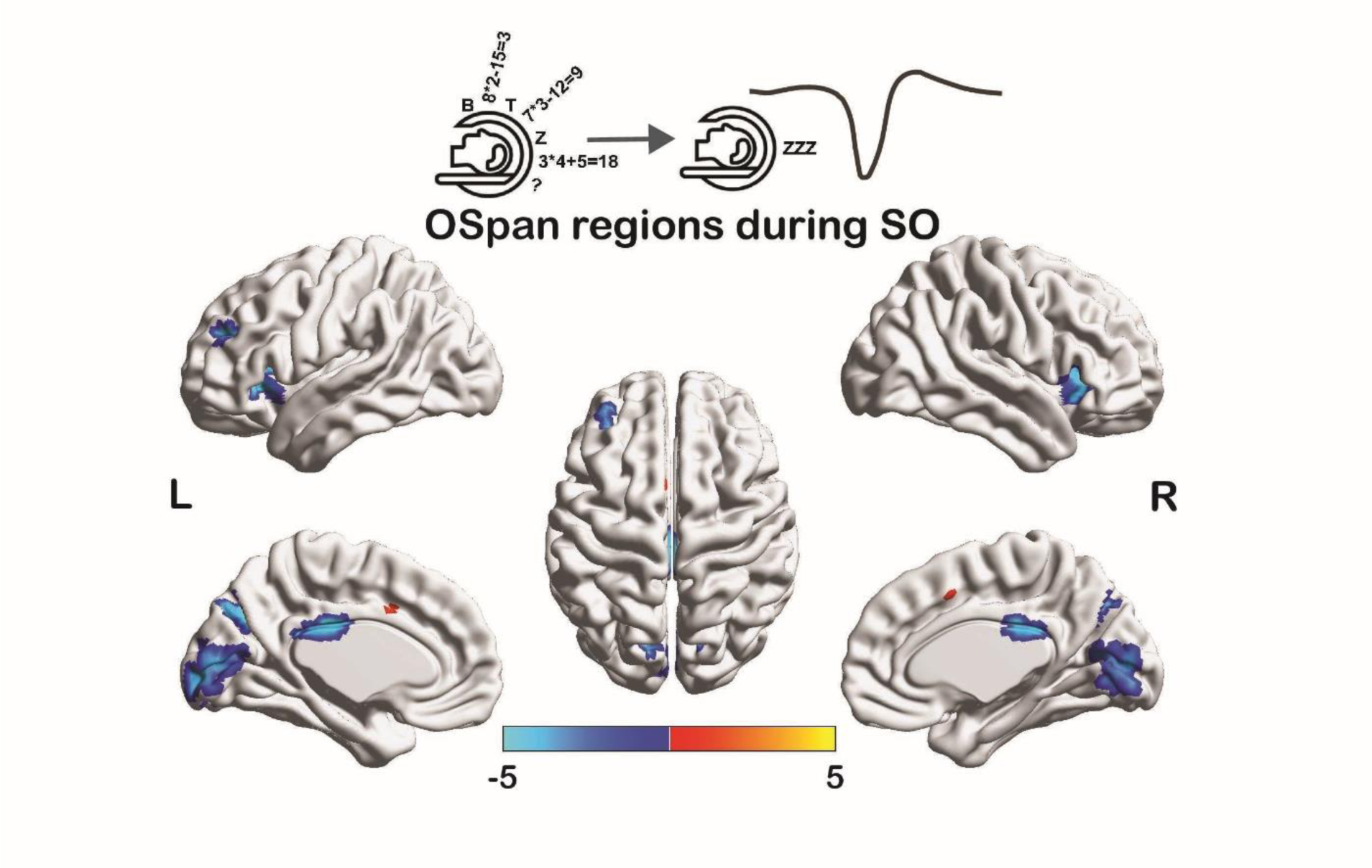
The activation of WM regions during SO windows. Warm color indicates higher activation during SO windows compared to non-SO NREM periods, and cold color indicates reduced activation.

### Correlations between Overnight OSPAN Changes and SO Activity

To assess whether changes in overnight WM activity may be related to neural processing during SOs, we examined correlations between these events using two distinct approaches. First, we asked whether variance in overnight (AM – PM) OSPAN (vs. Math) activation changes was related to variance in SO activation across individual sessions (Figure 5). This analysis was performed at the voxel-wise level, using robust linear regression (the robustfit function in MATLAB)^58^, which automatically downweights sessions with high variance (relative to the population, i.e. outliers). In other words, this analysis tested whether greater overnight changes in WM-related activation in a given brain region were associated with greater activation during SO events. Multiple comparisons were corrected using the Benjamini and Hochberg method with a false discovery rate q < 0.05 (significant clusters with an extent of > 5 voxels were reported). The scatterplot in Figure 5 illustrates the relationship between SO activation and overnight OSPAN (vs Math) activation change across each night (session in the study), averaged across all significant clusters.

**Figure 5.**
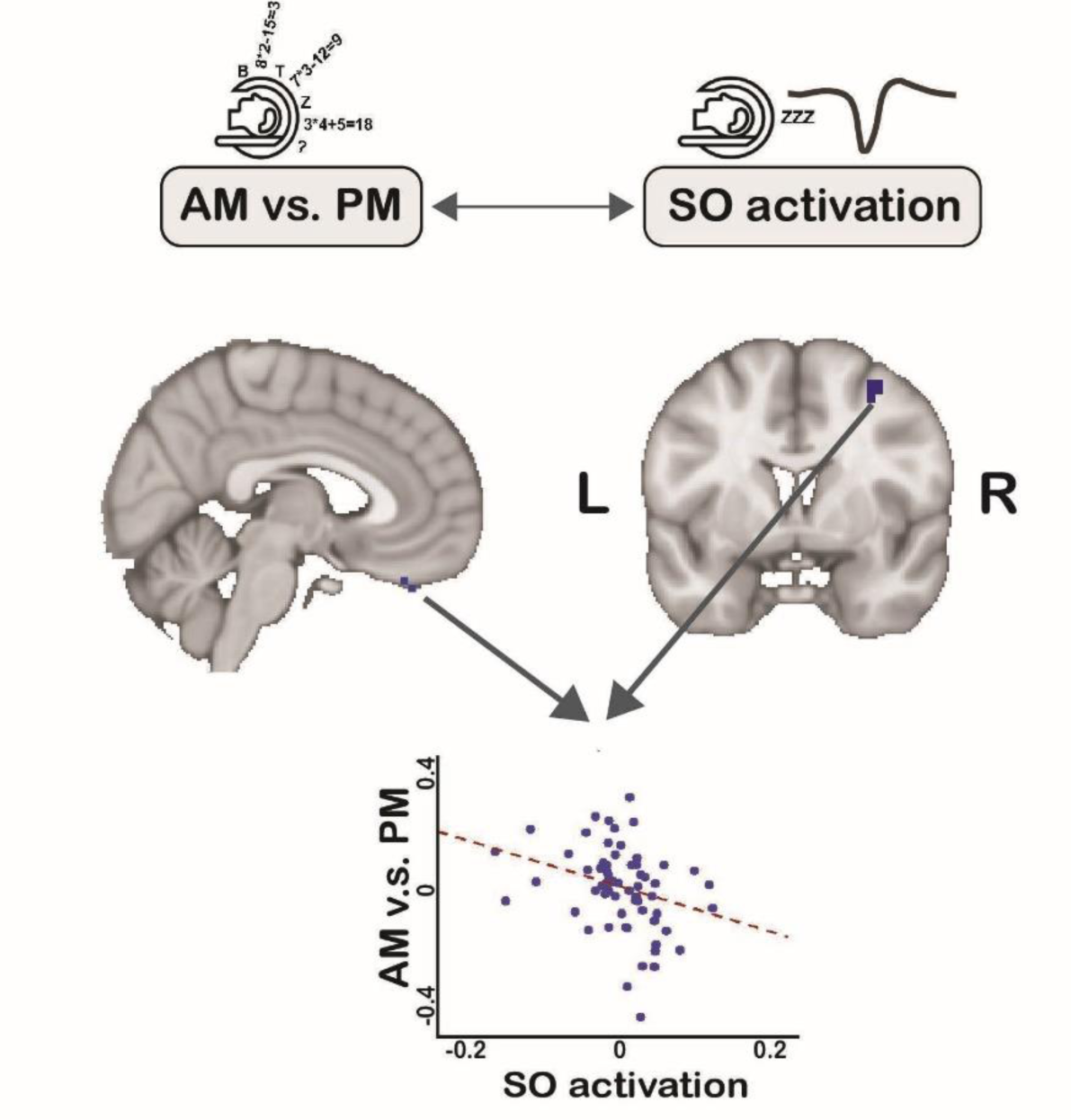
Correlation between overnight WM activation changes and SO activation, the two significant clusters are ventral medial prefrontal cortex (vmPFC) and premotor cortex.

In addition to the data-driven approach described above, we also used a network-driven approach to examine relationships between overnight changes in WM activity and SO activity. The goal of this analysis was to examine whether variance in overnight WM activation changes across voxels were related to the extent of activation during SOs in those voxels. To do so, we focused on pre-defined networks which have been implicated in attentional control. Specifically, we examined the Ventral Attention Network and Dorsal Attention Network, defined using the 7-network parcellation of the human cerebral cortex from Yeo et al. (2011)^59^. The Ventral Attention Network is shown on Figure 6a. To examine this relationship, we extracted the AM – PM change in activation (contrast value) and SO vs NREM activation (contrast value) and averaged these values across participants. We then performed robust linear regression to assess the correlation between overnight WM activity changes and SO activation across voxels (Figure 6c). The statistical significance of the observed correlation was assessed using a non-parametric permutation test. Specifically, we created a null distribution of correlation values by repeatedly shuffling the labels between the AM and PM sessions and re-computing the correlation between overnight changes and SO activation (N = 1000 permutations). Statistical significance (p-value shown in Figure 6c) was computed by comparing the observed correlation value to the null distribution.

**Figure 6.**
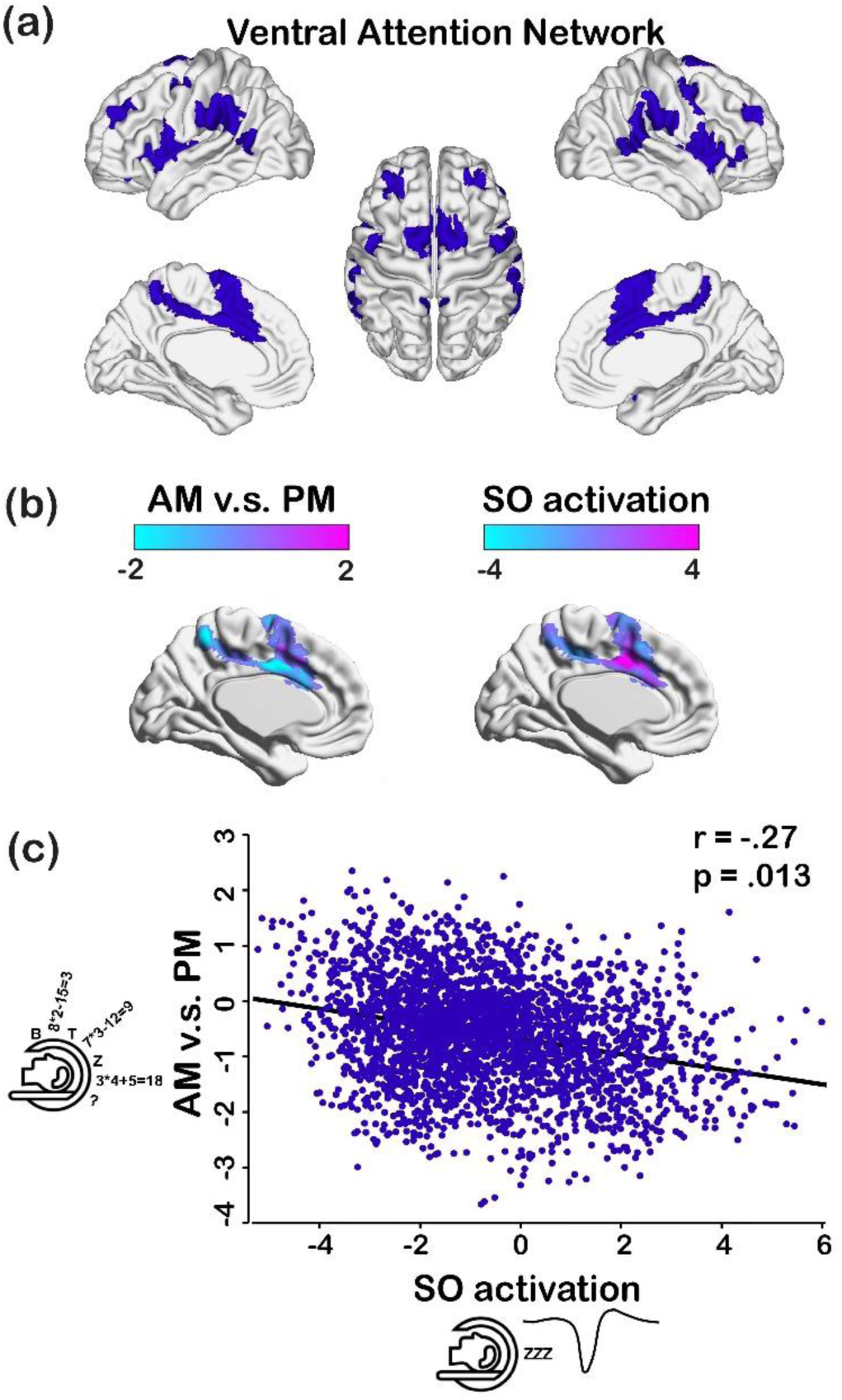
(a) The ventral attention network. (b) left: The overnight (AM-PM) activation change during OSPAN vs Math in the ventral attention network. right: The activation during SO windows compared to non-SO NREM windows in the ventral attention network. (c) Correlation between the AM-PM OSPAN activation change and SO activation.

## Results

### Sleep Architecture

On average, participants had 95.67 min (SD=36.62) of total sleep time in the MRI scanner (Table 1). The average sleep latency was 11.34 min (SD=12.73). The average duration of N2 sleep was 46.15 min (SD=22.59) and 36.3 min (SD=26.24) for SWS. NREM sleep from 34 participants (67 nights) was included in the analyses (for individual sleep architecture, see Supplemental Table 2). Throughout the night (sleep in the scanner + subsequent sleep outside of the scanner), participants were in bed for an average of 493.24 min (SD=73.78), with a total sleep time of 366.06 min (SD=108.39). OURA categorizes NREM sleep into Light and Deep sleep, which we combined with N2 and N3, respectively (Table 2).

### Behavioral Results

A 2-way repeated measures ANOVA (am/pm * visit 1/visit2) was used to assess changes in WM performance across sleep. Considering WM accuracy, significant main effects of time of day (F_(1,33)_=4.52, *p*=0.04) and visit (F_(1,33)_=10.94, *p*=0.002) were found, but no interaction between the two F(1,33)=1.92, p=0.18). The main effect of time of day was driven by significantly better WM accuracy during the AM test (M=0.77±0.18) compared to the PM test (M=0.74±0.16) (t_33_=-2.21, *p*=0.03; Figure 1c), demonstrating an overnight improvement in WM. Since no difference in overnight WM accuracy change was found across visits (no visit by time of day interaction), subsequent analyses measuring overnight changes in WM processing were performed on the data collapsed (averaged) across the two visits.

For RT, a 2-way repeated measures ANOVA (am/pm * visit 1/visit2) resulted in significant main effects of time of day (F_(1,33)_=5.89, *p*=0.02) and visit (F_(1,33)_=8.52, *p*=0.002), with a marginal interaction between the two F_(1,33)_=4.31, p=0.05). Participants had significantly faster responses in the AM (M=1.19±0.31 s) compared to PM (M=1.26±0.29 s) for all correct OSPAN trials (t_33_=2.43, *p*=0.02). Together, these results demonstrate that WM function improved across sleep in our study, both in faster and more accurate responses, as expected from prior work^7,8,32^.

### Neuroimaging Results

#### 1) Working Memory-related Activation

We first asked whether changes in WM function across sleep were reliable in our dataset. To do so, we assessed whether there is a reliable or similar pattern of overnight changes in WM-related activation (difference between AM and PM sessions for the OSPAN vs. Math contrast, see Methods) across the two visits within each participant. Therefore, we measured the whole-brain voxel-wise correlation of overnight activation change between the two visits in each participant. We found that the correlation in overnight WM-related activity change was significantly greater from zero (AM – PM; Mean r=0.06, t=2.34, *p*=0.03). This suggests that the patterns of WM-related activation change between AM and PM sessions were stable across visits.

Before isolating regions that showed specific overnight changes in WM activation, we characterized general activation associated with WM in our dataset (WM-related regions). To do so, we examined the OSPAN vs. Math contrast at the whole-brain level across all sessions. Greater activation during OSPAN compared to Math blocks was present in several cortical and subcortical areas (Figure 2a warm color), including the bilateral precentral gyrus, primary motor cortex, dorsal anterior cingulate cortex, caudate, anterior insular cortex, thalamus, putamen, occipital cortex and dorsal precuneus. Conversely, activation was reduced during OSPAN compared to Math blocks in dorsal medial prefrontal cortex (PFC), anterior frontal, inferior parietal areas, and left lateral temporal cortex (Figure 2a cold color, FWE corrected at *p*<0.05).

To assess whether WM-related activation showed reliable overnight changes, or differed between PM and AM sessions, we first characterized WM activation in each PM and AM session separately. During the PM session, we found widespread activation during OSPAN vs Math blocks encompassing the frontal and parietal cortices (Figure 2b blue, FWE corrected at *p*<0.05), including the lateral PFC, premotor cortex, dorsal precuneus, dorsal anterior cingulate cortex, anterior insular cortex, putamen, caudate, occipital cortex, and supramarginal gyrus. Overall, less WM-related activation was present during the AM session (Figure 2b red, FWE corrected at *p*<0.05), including regions in the frontal cortex (premotor and motor cortex and inferior frontal gyrus), and subcortical regions including cingulate and left putamen. These results qualitatively suggest that fewer brain regions were reliably recruited during WM during AM compared to PM sessions.

To determine whether WM activation was reliably reduced across sleep, we assessed the change in OSPAN vs Math activation between the AM and PM sessions within regions engaged during WM (shown in Fig 2a warm color). The dorsal precuneus region (MNI: 8.3, -59, 53.4) showed a significant decrease in activation overnight (Figure 2c; FWE corrected at *p*<0.05). We further defined this significant cluster in the precuneus as our target ROI (seed region) for subsequent PPI analyses.

To summarize, we found widespread activation during the PM session, while less activation occurred during the AM session, suggesting fewer brain regions were recruited for WM after sleep. We then revealed a significant decrease in activation overnight in the dorsal precuneus.

#### 2) Functional connectivity of the dorsal precuneus (target ROI)

To investigate the functional interactions of the dorsal precuneus, the WM region significantly modulated by sleep, we examined whole-brain functional connectivity using PPI with target precuneus ROI during OSPAN vs. Math blocks (Figure 3a, FWE corrected at *p*<0.05). WM-related connectivity of the dorsal precuneus during OSPAN vs Math blocks was found with multiple cortical sites, including the fusiform gyrus, superior parietal cortex, middle frontal gyrus, and premotor cortex. Reduced connectivity during OSPAN vs. Math blocks was found between the dorsal precuneus and small regions in occipital and inferior parietal cortex (Figure 3a, FWE-corrected at *p*<0.05).

To test whether SOs modulate brain-wide functional connectivity of the dorsal precuneus, we also examined connectivity of the target ROI during SOs during NREM sleep (the SO vs non-SO NREM contrast, see Methods) (Figure 3b). The dorsal precuneus showed a diffuse decrease in functional connectivity with frontal and parietal cortices during SOs relative to non-SO NREM sleep (FWE-corrected at *p*<0.05). This result suggests a role for SOs in modulating the whole-brain connectivity between areas showing sleep-dependent change in WM activation.

#### 3) Working memory BOLD Activation during SO

To better understand the relation between neural processing during WM and SOs, we examined if the WM-related regions that we isolated (shown in Figure 2a warm color) were reliably modulated by SOs. We observed BOLD activation in WM-related regions during SO events compared to non-SO NREM. Specifically, portions of the dorsal mid-cingulate cortex and right thalamus were activated during SOs (Figure 4 warm color, FWE corrected at *p*<0.05), and decreased activation (relative to non-SO NREM) was found in the anterior insular cortex, putamen, dorsal precuneus and occipital cortex (Figure 4 cold color, FWE corrected at *p*<0.05). These data suggest that WM-related regions were modulated during SOs, which further suggests the question of how online task-related activity might be reflected during SO windows.

#### 4) Correlation between working memory and SO BOLD activation across sessions

To establish a link between overnight changes in WM function and neural processing during SOs, we asked whether overnight WM activation changes were related to the extent of SO activation across sessions. We found a negative correlation between overnight WM changes (AM – PM) and activation during SOs across sessions in the ventral medial prefrontal cortex (vmPFC) and premotor cortex (Figure 5, FWE-corrected at *p*<0.05, cluster extent threshold > 5 voxels). In other words, greater activity in the vmPFC and premotor cortex during SOs was associated with reduced activity during online WM (OSPAN vs Math blocks) after a night of sleep, suggesting a re-organization of frontal WM-related activity during SOs.

#### 5) Spatial correlation between working memory and SO BOLD activation in the Ventral Attention Network

To investigate the potential role of neural processing during SOs in modulating overnight changes in WM function, we also examined pre-defined networks implicated in cognitive control, the dorsal attention and ventral attention (salience) networks (Figure 6a). We first tested whether these networks showed significant WM-related activity in our task (greater activation during OSPAN vs. Math blocks). Indeed, the ventral attention network was more active during OSPAN vs. Math blocks, (t=3.02, *p*=0.0045), while the activation in the dorsal attention network did not differ between OSPAN and Math blocks (t=1.62 *p*=0.11).

To test whether activity during SOs may modulate overnight changes in WM activity, we examined whether patterns of overnight (AM-PM) WM (OSPAN vs. Math) activation changes across voxels were related to activation patterns during SOs (vs. non-SO NREM, Figure 6b). Across voxels in the ventral attention network, there was a negative correlation between patterns of overnight OSPAN activation change and SO activation (r=-0.27, *p*=0.01, N=1000 permutations). This indicates that voxels with greater overnight reductions in WM activation show stronger activation during SO events. This finding suggests that ventral attention network activity during SOs might facilitate increased reduction of WM activation across a night of sleep. This pattern was not found in the dorsal attention network, as the correlation between patterns of overnight OSPAN activation change and SO activation was not statistically significant across voxels in this network (r=-0.11, *p*=0.25, N=1000 permutations).

## Discussion

We investigated the neural basis of WM improvement during sleep using simultaneous EEG-fMRI. To our knowledge, this is the first study to demonstrate how sleep events, specifically SOs, may orchestrate functional changes in the brain associated with WM improvement. Our findings reveal several important insights into how the neural mechanisms supporting WM change over a night of sleep. First, we identified the effects of an overnight sleep on WM-related BOLD activation. Overnight sleep is associated with reliable changes in WM activity across sessions and specifically, decreased recruitment of the dorsal precuneus. We demonstrated that SOs modulated BOLD activation in WM-related regions. Critically, we uncovered multiple relationships between the extent of neural processing during SOs and changes in overnight WM activity. Across sessions, stronger activation in the vmPFC and premotor cortex during SOs was associated with greater overnight reductions in WM activity. Additionally, multivoxel patterns of activation during SOs were similar to patterns of overnight reductions in WM activity within the Ventral Attention Network, suggesting that SO activity may modulate changes in WM function. Together, these results provide evidence for a direct link between SO activity and overnight changes in WM function, which may underlie improvements WM across sleep.

Here, we found reduced recruitment of the dorsal precuneus during WM across a night of sleep. Prior fMRI studies of online WM processing have consistently implicated the posterior parietal cortex^1,60,61^ and the precuneus specifically^62–64^ in WM function. For example, one study showed greater activation in the precuneus during a spatial WM task compared to other structures in the posterior parietal region^65^. In a meta-analysis of 11 fMRI studies using a n-back WM paradigm, the precuneus was highlighted^66^, along with an MEG study using a similar task^67^. Another meta-analysis also indicated precuneus in the WM process, and further proposed that ventral frontal and temporal regions mediate object instead of spatial WM^40^. Moreover, the precuneus has been shown to reflect the cognitive demands of WM, as increases in task difficulty on an N-back task (i.e., slower RT) was associated with greater activation in the precuneus^60^. Studies have established a direct role of the precuneus in WM via transcranial magnetic stimulation (TMS). Specifically, TMS applied to the midline parietal site centered around the precuneus led to improved WM performance, whereas TMS applied to other regions such as frontal or occipital sites did not have such benefit^68^. Improved WM performance has been associated with distinct changes in the connectivity between the precuneus and other brain networks. A prior study reported improved WM performance after a 6-week WM training is associated with reduced functional connectivity in the default mode network, including the precuneus, and increased functional connectivity in the frontoparietal network^69^. Building on these findings, we show that the recruitment of the precuneus was reduced after sleep, in parallel with improvements in WM function across sleep, suggesting that the precuneus may play a role in sleep-dependent WM improvement.

In addition to WM function, the precuneus has been implicated in long-term memory consolidation. Researchers have proposed that the precuneus, as part of the medial posterior parietal cortex, may constitute a memory engram for specific tasks^70^. Memory engram represents the specific group of neurons or neural connections that are associated with a particular memory^71,72^. Specifically, using an object-location declarative memory task, researchers detected significant learning-induced microstructural changes and multivoxel stimulus representations in the precuneus, which correlated with better performance^73^. While our WM paradigm substantially differs from the learning of specific object-location memories, it is possible to speculate that changes in precuneus activity could reflect the development of a task-specific representation, e.g., memory of the specific task procedures and strategies, or monitoring and maintenance of visual letter representations. In other words, improvements in WM function and changes in precuneus activity could potentially reflect consolidating a memory of the task as a whole, which may allow more resources to be used for WM capacity. This hypothetical task-specific learning process is congruent with the inconsistent transfer of learning on one WM task to other untrained executive function tasks^74–76^, and is consistent with the idea that transfer may occur when tasks share that same underlying principles or knowledge structure^77^. However, we did not test this idea directly in the current study, and the possibility that precuneus activity reflects a task-specific representation would need to be explored in future work.

After showing reduced precuneus activity over a night of sleep, in parallel with WM improvement, the next logical question is: What are the specific neural mechanisms during sleep that led to reduced WM-related activation? To this end, our findings suggest that NREM SOs may play a role in modulating WM activity. During SOs, we found a broad decrease in activation in WM-related areas and a decrease in precuneus connectivity with other WM-related regions. The decrease in precuneus connectivity may potentially reflect decoupling of the precuneus from WM-related networks. Furthermore, reduced neural activation during SOs is consistent with prior findings showing a broad decrease in activation in frontal and parietal areas as well as subcortical areas including the precuneus during NREM sleep using fMRI^78^ and PET^26^. SWS has been thought of as a state of quiescence, during which brain oxygen metabolism, cerebral blood flow, and glucose metabolism are decreased compared to wakefulness and REM sleep^79–81^. However, SOs periods are not a total silence of communication. We also found increased activation in the cingulate and right thalamus during SOs compared to non-SO NREM periods, suggesting SOs modulate WM-related network by selectively activating specific areas while silencing others. Indeed, we found that the more WM-related areas were active during SOs, the less they were activated after sleep compared to pre-sleep, suggesting that fewer online resources may be needed when regions are active during SOs. Given the established role of SOs in contributing to neural communication supporting long-term memory^82^, together with our results, SOs may provide the temporal framework for systems level communication which may be important for both WM and episodic memory processing.

Importantly, when probing direct associations between brain activity during SOs and during WM processing, we found that when WM-related areas are active during SOs, they seem to require fewer resources compared to before sleep. We observed this negative correlation between SOs-related activation and task-related activation using two methods. First, we found a negative correlation between the extent of SO activity across sessions and reductions in WM activity in the premotor cortex and vmPFC, both regions that were engaged in WM processing in our dataset. Moreover, multivoxel activation patterns during SOs were similar, or tended to mirror patterns of overnight changes in WM function in the Ventral Attention Network, such that voxels with greater activity during SOs showed larger reductions in overnight WM activity. These results show explicit links between SOs and overnight changes in WM function during online processing.

The specific neural mechanism underlying the role of SOs in modulating WM-networks remains to be explored. One the one hand, studies have shown that SOs in SWS promote synaptic downscaling, which involves the weakening of synaptic connections that have been strongly activated during wakefulness^21^. This downscaling process is thought to help maintain the stability and efficiency of neural networks, preventing them from becoming overloaded and potentially facilitate cognitive functioning including WM by freeing up capacity for the encoding of new information during the upcoming wake period^83^. Another potential mechanism of how SOs modulate WM-networks is through glymphatic clearance. The glymphatic system is responsible for brain-wide delivery of nutrients and clearance of waste via influx of cerebrospinal fluid (CSF) alongside perivascular spaces and through the brain^84^. Specifically, SWS is associated with glymphatic clearance, as studies have reported that SOs are temporally coupled with and precede cerebrospinal glymphatic clearance^85,86^. Thus, an indirect benefit of clearing metabolic waste products and toxic substances that accumulate in the brain might be to maintain the optimal functioning of the brain, including the networks involved in WM. Further research that elucidates multi-level brain activity during SOs across animal and humans is needed to better understand the mechanism underlying SOs’ modulation in WM-networks.

Our study has several limitations. The limited sleep duration and the amount of recorded sleep events during the scan made it impossible to differentiate between SOs in N2 and SWS, as SOs show different characteristics between the two sleep stages^87^. It remains to be explored how SOs modulate WM-related areas throughout the night as we were only able to collect EEG-fMRI data for the first sleep cycle^88,89^. Moreover, our study focuses on WM-related functional changes during sleep. Previous studies have shown WM and episodic memory might compete for limited resources during sleep, and particularly SOs^90^. Therefore, it will be interesting to investigate how neural processing during SOs may reflect this competition between WM and episodic memory.

In conclusion, our findings suggest that overnight sleep plays a critical role in WM by modulating neural processing during WM. By showing multiple links between neural processing during SOs and overnight changes in WM function, our study highlights the potential importance of SOs in modulating neural activity supporting WM. These findings provide a basis to develop theoretic models on sleep-dependent WM benefit. It may also have implications for the development of sleep-based interventions aimed at improving WM performance.

