## Supplementary material for "Slow Oscillations Modulate Functional Brain Changes Supporting Working Memory": suppl

***<u>Supplemental Results</u>***

### *Table 1. Time in bed in the scanner*

#TR of the sleep scans that contain NREM sleep

| | | Visit 1 | Visit 2 |
|---|---|---|---|
| 1 | 105 | no NREM | 4096 |
| 2 | 106 | 1036+824+1735 | 2995 |
| 3 | 107 | 4095 | 2024+1841 |
| 4 | 108 | 2113+ 1001 | 4096 |
| 5 | 111 | 4006 | 4096 |
| 6 | 112 | 856+1565 | 1598 |
| 7 | 113 | 4096 | 4096 |
| 8 | 114 | 2734 | 4096 |
| 9 | 115 | 3703 | 4096 |
| 10 | 116 | 1246+2192 | 3643 |
| 11 | 118 | 2108+2093 | 2887 |
| 12 | 121 | 2327 | 3349 |
| 13 | 123 | 1478 | 2812+1843 |
| 14 | 124 | 4096 | 2297+2085 |
| 15 | 125 | 3648 | 3436 |
| 16 | 126 | 3261 | 3458 |
| 17 | 127 | 4096 | 3887 |
| 18 | 130 | 1184+4094 | 4096 |
| 19 | 131 | 3448 | No NREM |
| 20 | 132 | 4096 | 4096 |
| 21 | 133 | 4040 | 3093 |
| 22 | 135 | 4096 | 3862 |
| 23 | 136 | 3688 | 3631 |
| 24 | 137 | 2612+1474 | 3447 |
| 25 | 138 | 3688 | 2504 |
| 26 | 142 | 3647 | 3782 |
| 27 | 144 | 4096 | 4096 |
| 28 | 148 | 4096 | 4096 |
| 29 | 149 | 4096 | 3775 |
| 30 | 152 | 3345 | 1686+911 |
| 31 | 153 | 3726 | 3397 |
| 32 | 154 | 4096 | 4096 |
| 33 | 156 | 4096 | 4096 |
| 34 | 160 | 4096 | 4096 |

### *Table 2. Sleep architecture in the scanner*

| SID | Visit | TIB | TST | s1Min | s2Min | s3Min | remMin | sleepOnSet | s2OnSet | s3OnSet | remOnSet | WASO | SE |
|---|---|---|---|---|---|---|---|---|---|---|---|---|---|
| 105 | 2 | 56.5 | 44 | 15 | 26 | 3 | 0 | 1.5 | 16.5 | 34.5 | 0 | 0 | 77.88 |
| 106 | 1 | 107.5 | 42 | 14 | 21 | 7 | 0 | 6 | | | | 44.5 | 39.07 |
| 106 | 2 | 111.5 | 39 | 25 | 14 | 0 | 0 | 18 | 23 | 0 | 0 | 27.5 | 34.98 |
| 107 | 1 | 151.5 | 150 | 2.5 | 65.5 | 82 | 0 | 1 | 3 | 10 | 0 | 1 | 99.00990099 |
| 107 | 2 | 143.5 | 142.5 | 0.5 | 54 | 69.5 | 18.5 | 1 | | | | 1 | 99.30313589 |
| 108 | 1 | 120 | 88.5 | 32.5 | 47.5 | 8.5 | 0 | 4 | | | | 17 | 73.75 |
| 108 | 2 | 152.5 | 108.5 | 48 | 54 | 6.5 | 0 | 8.5 | 17 | 43 | 0 | 36 | 71.14754098 |
| 111 | 1 | 150 | 140.5 | 2.5 | 79.5 | 55.5 | 0 | 7 | 9.5 | 39 | 0 | 3 | 93.66666667 |
| 111 | 2 | 79 | 65.5 | 5 | 32.5 | 28 | 0 | 12 | 15.5 | 29 | 0 | 1 | 82.91139241 |
| 112 | 1 | 89.5 | 55 | 2 | 33.5 | 19.5 | 0 | 8.5 | | | | 22 | 61.45251397 |
| 112 | 2 | 62 | 44.5 | 0 | 26.5 | 18 | 0 | 9 | 9 | 35.5 | 0 | 0 | 71.77419355 |
| 113 | 1 | 153 | 138 | 13 | 100.5 | 24.5 | 0 | 1.5 | 4 | 80.5 | 0 | 14 | 90.19607843 |
| 113 | 2 | 153 | 148.5 | 1 | 62 | 82.5 | 0 | 1 | 1 | 6 | 0 | 0 | 97.05882353 |
| 114 | 1 | 102.5 | 78.5 | 4 | 51 | 23.5 | 0 | 16 | 20 | 39.5 | 0 | 0 | 76.58536585 |
| 114 | 2 | 153 | 41 | 20.5 | 20.5 | 0 | 0 | 19.5 | 54.5 | 0 | 0 | 74.5 | 26.79738562 |
| 115 | 1 | 138 | 71.5 | 7.5 | 33.5 | 30 | 0 | 12 | 18.5 | 23.5 | 0 | 2 | 51.8115942 |
| 115 | 2 | 153 | 114.5 | 4 | 65.5 | 43.5 | 0 | 9.5 | 11 | 20 | 0 | 12 | 74.83660131 |
| 116 | 1 | 230.5 | 102 | 20 | 66.5 | 12.5 | 0 | 20.5 | | | | 83 | 44.2516269 |
| 116 | 2 | 149 | 56.5 | 12.5 | 44 | 0 | 0 | 9.5 | | | | 66.5 | 37.91946309 |
| 118 | 1 | 157 | 87.5 | 30.5 | 41 | 16 | 0 | 7.5 | | | | 56 | 55.73248408 |
| 118 | 2 | 108.5 | 54 | 10.5 | 39.5 | 4 | 0 | 30.5 | 36.5 | 48.5 | 0 | 4.5 | 49.76958525 |
| 121 | 1 | 50.5 | 32 | 2 | 2 | 0 | 28 | 3 | 30 | 0 | 16 | 16 | 63.36633663 |
| 121 | 2 | 125 | 73 | 20 | 13 | 0 | 40 | 48.5 | 108 | 0 | 67 | 4 | 58.4 |
| 123 | 1 | 56 | 52 | 0 | 13.5 | 38.5 | 0 | 0.5 | 0.5 | 13.5 | 0 | 0 | 92.85714286 |
| 123 | 2 | 97 | 78.5 | 5 | 23.5 | 50 | 0 | 12.5 | | | | 7 | 80.92783505 |
| 124 | 1 | 153 | 151.5 | 1 | 62 | 87.5 | 1 | 2 | 3 | 19 | 92 | 0 | 99.01960784 |
| 124 | 2 | 163.5 | 135.5 | 9 | 73.5 | 53 | 0 | 5.5 | | | | 16.5 | 82.87461774 |
| 125 | 1 | 137 | 66.5 | 28 | 37 | 1.5 | 0 | 48.5 | 81 | 97.5 | 0 | 9.5 | 48.54014599 |
| 125 | 2 | 129 | 95 | 19 | 50 | 21 | 0 | 2.5 | 7 | 58 | 0 | 3.5 | 73.64341085 |
| 126 | 1 | 121.5 | 63.5 | 5.5 | 20 | 37 | 0 | 10 | 12 | 20 | 0 | 19.5 | 52.26337449 |
| 126 | 2 | 129.5 | 48 | 1 | 21 | 26 | 0 | 2.5 | 3.5 | 13.5 | 0 | 0 | 37.06563707 |
| 127 | 1 | 152.5 | 141 | 6 | 97 | 36.5 | 0 | 0.5 | 1.5 | 6 | 0 | 11.5 | 92.45901639 |
| 127 | 2 | 145.5 | 93.5 | 16.5 | 39 | 38 | 0 | 1 | 1.5 | 9.5 | 0 | 14.5 | 64.26116838 |
| 130 | 1 | 197 | 41 | 7 | 21 | 13 | 0 | 14.5 | | | | 129.5 | 20.81218274 |
| 130 | 2 | 153 | 97.5 | 1 | 57.5 | 38 | 0 | 3.5 | 4 | 19.5 | 0 | 52.5 | 63.7254902 |
| 131 | 1 | 128.5 | 98.5 | 9 | 70 | 19.5 | 0 | 10 | 11 | 21 | 0 | 20.5 | 76.6536965 |
| 131 | 2 | 262.5 | 141.5 | 73.5 | 64 | 0 | 2 | 24 | | | | 32 | 53.9047619 |
| 132 | 1 | 153.5 | 115.5 | 13.5 | 75 | 27 | 0 | 7.5 | 11 | 20.5 | 0 | 19.5 | 75.24429967 |
| 132 | 2 | 184.5 | 24 | 7.5 | 4 | 12.5 | 0 | 10.5 | 18 | 20.5 | 0 | 0 | 13.00813008 |

| | | | | | | | | | | | | | |
|---|---|---|---|---|---|---|---|---|---|---|---|---|---|
| 133 | 1 | 151 | 90 | 0.5 | 52.5 | 37 | 0 | 1 | 1.5 | 2.5 | 0 | 0 | 59.60264901 |
| 133 | 2 | 116 | 104 | 1 | 42.5 | 60.5 | 0 | 7.5 | 8.5 | 20.5 | 0 | 0 | 89.65517241 |
| 135 | 1 | 153 | 148 | 6.5 | 85.5 | 56 | 0 | 5.5 | 10 | 24.5 | 0 | 0 | 96.73202614 |
| 135 | 2 | 144.5 | 138.5 | 16.5 | 56.5 | 65.5 | 0 | 1.5 | 15.5 | 26.5 | 0 | 5 | 95.84775087 |
| 136 | 1 | 137.5 | 134 | 3 | 67.5 | 60 | 0 | 3.5 | 4 | 16 | 0 | 0 | 97.45454545 |
| 136 | 2 | 136 | 135 | 0 | 48 | 59 | 23.5 | 1 | 1 | 2 | 90.5 | 0 | 99.26470588 |
| 137 | 1 | 152 | 81 | 9 | 59.5 | 12 | 0 | 11.5 | | | | 23.5 | 53.28947368 |
| 137 | 2 | 128.5 | 72 | 6.5 | 48.5 | 16.5 | 0 | 4.5 | 11 | 36.5 | 0 | 42.5 | 56.0311284 |
| 138 | 1 | 138 | 85.5 | 2 | 62.5 | 21 | 0 | 21.5 | 22.5 | 32 | 0 | 26.5 | 61.95652174 |
| 138 | 2 | 93.5 | 76 | 2 | 74 | 0 | 0 | 3 | 3 | 0 | 0 | 13 | 81.28342246 |
| 142 | 1 | 143 | 134 | 6 | 53 | 75 | 0 | 1.5 | 7.5 | 15 | 0 | 0 | 93.70629371 |
| 142 | 2 | 141 | 124.5 | 45 | 23 | 56.5 | 0 | 7 | 62 | 74.5 | 0 | 10 | 88.29787234 |
| 144 | 1 | 331.5 | 131 | 14 | 53.5 | 50 | 12.5 | 15 | | | | 43 | 39.5173454 |
| 144 | 2 | 153 | 150.5 | 1 | 84.5 | 61.5 | 0 | 2.5 | 3.5 | 10.5 | 0 | 0 | 98.36601307 |
| 148 | 1 | 152.5 | 127 | 17.5 | 64 | 45.5 | 0 | 0.5 | 0.5 | 5 | 0 | 25.5 | 83.27868852 |
| 148 | 2 | 153 | 123 | 21 | 47.5 | 54 | 0 | 21 | 26.5 | 33 | 0 | 9.5 | 80.39215686 |
| 149 | 1 | 153 | 105 | 8.5 | 58.5 | 36.5 | 0 | 5 | 7.5 | 23.5 | 0 | 43.5 | 68.62745098 |
| 149 | 2 | 141.5 | 135.5 | 7.5 | 52 | 75 | 0 | 1 | 2 | 11 | 0 | 5 | 95.75971731 |
| 152 | 1 | 125 | 109 | 4 | 24.5 | 80.5 | 0 | 15 | 19 | 21.5 | 0 | 1 | 87.2 |
| 152 | 2 | 97.5 | 48 | 7.5 | 14.5 | 26 | 0 | 50 | 57.5 | 37.5 | 0 | 0 | 49.23076923 |
| 153 | 1 | 139 | 93 | 1.5 | 46 | 40 | 0 | 1.5 | 3 | 12.5 | 0 | 0.5 | 66.90647482 |
| 153 | 2 | 126.5 | 108 | 5 | 53.5 | 48 | 0 | 2.5 | 4 | 13 | 0 | 5.5 | 85.37549407 |
| 154 | 1 | 153 | 132 | 6 | 23.5 | 102 | 0 | 14.5 | 19.5 | 27 | 0 | 7 | 86.2745098 |
| 154 | 2 | 153 | 41 | 6.5 | 7.5 | 27 | 0 | 16 | 21 | 28.5 | 0 | 1.5 | 26.79738562 |
| 156 | 1 | 153 | 117 | 6.5 | 66 | 44.5 | 0 | 19 | 20 | 29.5 | 0 | 3.5 | 76.47058824 |
| 156 | 2 | 153 | 96 | 1.5 | 31.5 | 63 | 0 | 56.5 | 57.5 | 66 | 0 | 1 | 62.74509804 |
| 160 | 1 | 153 | 116 | 7.5 | 39.5 | 69 | 0 | 26 | 28.5 | 56 | 0 | 11.5 | 75.81699346 |
| 160 | 2 | 129 | 95.5 | 4.5 | 31.5 | 57.5 | 0 | 34 | 38.5 | 49 | 0 | 0 | 74.03100775 |

***Table 3. Detected SOs***

| SID | Visit | SO count | SO density |
|---|---|---|---|
| 105 | 2 | 92 | 3.17 |
| 106 | 1 | 136 | 4.69 |
| 106 | 2 | 57 | 1.97 |
| 107 | 1 | 901 | 31.07 |
| 107 | 2 | 743 | 25.62 |
| 108 | 1 | 143 | 4.93 |
| 108 | 2 | 275 | 9.48 |
| 111 | 1 | 609 | 21.00 |

| | | | |
|---|---|---|---|
| 111 | 2 | 259 | 8.93 |
| 112 | 1 | 198 | 6.83 |
| 112 | 2 | 203 | 7.00 |
| 113 | 1 | 697 | 24.03 |
| 113 | 2 | 820 | 28.28 |
| 114 | 1 | 429 | 14.79 |
| 114 | 2 | 100 | 3.45 |
| 115 | 1 | 411 | 14.17 |
| 115 | 2 | 557 | 19.21 |
| 116 | 1 | 225 | 7.76 |
| 116 | 2 | 168 | 5.79 |
| 118 | 1 | 316 | 10.90 |
| 118 | 2 | 239 | 8.24 |
| 121 | 2 | 82 | 2.83 |
| 123 | 1 | 130 | 4.48 |
| 123 | 2 | 422 | 14.55 |
| 124 | 1 | 761 | 26.24 |
| 124 | 2 | 642 | 22.14 |
| 125 | 1 | 163 | 5.62 |
| 125 | 2 | 354 | 12.21 |
| 126 | 1 | 377 | 13.00 |
| 126 | 2 | 278 | 9.59 |
| 127 | 1 | 701 | 24.17 |
| 127 | 2 | 372 | 12.83 |
| 130 | 1 | 186 | 6.41 |
| 130 | 2 | 485 | 16.72 |
| 131 | 1 | 468 | 16.14 |
| 132 | 1 | 653 | 22.52 |
| 132 | 2 | 133 | 4.59 |
| 133 | 1 | 552 | 19.03 |
| 133 | 2 | 608 | 20.97 |
| 135 | 1 | 839 | 28.93 |
| 135 | 2 | 754 | 26.00 |
| 136 | 1 | 769 | 26.52 |
| 136 | 2 | 475 | 16.38 |
| 137 | 1 | 429 | 14.79 |
| 137 | 2 | 386 | 13.31 |
| 138 | 1 | 544 | 18.76 |
| 138 | 2 | 367 | 12.66 |
| 142 | 1 | 790 | 27.24 |

| | | | |
|---|---|---|---|
| 142 | 2 | 443 | 15.28 |
| 144 | 1 | 283 | 9.76 |
| 144 | 2 | 752 | 25.93 |
| 148 | 1 | 666 | 22.97 |
| 148 | 2 | 668 | 23.03 |
| 149 | 1 | 516 | 17.79 |
| 149 | 2 | 655 | 22.59 |
| 152 | 1 | 562 | 19.38 |
| 152 | 2 | 214 | 7.38 |
| 153 | 1 | 368 | 12.69 |
| 153 | 2 | 680 | 23.45 |
| 154 | 1 | 760 | 26.21 |
| 154 | 2 | 269 | 9.28 |
| 156 | 1 | 714 | 24.62 |
| 156 | 2 | 570 | 19.66 |
| 160 | 1 | 588 | 20.28 |
| 160 | 2 | 442 | 15.24 |

### ***Supplemental Methods***

#### *Anatomical data preprocessing*

A total of 6 T1-weighted (T1w) images were found within the input BIDS dataset. All of them were corrected for intensity non-uniformity (INU) with N4BiasFieldCorrection (Tustison et al. 2010), distributed with ANTs 2.2.0 (Avants et al. 2008, RRID:SCR_004757). The T1w-reference was then skull-stripped with a Nipype implementation of the antsBrainExtraction.sh workflow (from ANTs), using OASIS30ANTs as target template. Brain tissue segmentation of cerebrospinal fluid (CSF), white-matter (WM) and gray-matter (GM) was performed on the brain-extracted T1w using fast (FSL 5.0.9, RRID:SCR_002823, Zhang, Brady, and Smith 2001). A T1w-reference map was computed after registration of 6 T1w images (after INU-correction) using mri_robust_template (FreeSurfer 6.0.1, Reuter, Rosas, and Fischl 2010). Volume-based spatial normalization to one standard space (MNI152NLin2009cAsym) was performed through nonlinear registration with antsRegistration (ANTs 2.2.0), using brain-extracted versions of both T1w reference and the T1w template. The following template was selected for spatial normalization: ICBM 152 Nonlinear Asymmetrical template version 2009c [Fonov et al. (2009), RRID:SCR_008796; TemplateFlow ID: MNI152NLin2009cAsym].

#### *Functional data preprocessing*

For each of the BOLD runs found per subject (across all tasks and sessions), the following preprocessing was performed. First, a reference volume and its skull-stripped version were generated using a custom methodology of fMRIPrep. The BOLD reference was then co-registered to the T1w reference using flirt (FSL 5.0.9, Jenkinson and Smith 2001) with the boundary-based registration (Greve and Fischl 2009) cost-function. Co-registration was configured with nine degrees of freedom to account for distortions remaining in the BOLD reference. Head-motion parameters with respect to the BOLD reference (transformation matrices, and six corresponding rotation and translation parameters) are estimated before any spatiotemporal filtering using mcflirt (FSL 5.0.9, Jenkinson et al. 2002). BOLD runs were slice-time corrected using 3dTshift from AFNI 20160207 (Cox and Hyde 1997, RRID:SCR_005927). The BOLD time-series (including slice-timing correction when applied) were resampled onto their original, native space by applying a single, composite transform to correct for head-motion and susceptibility distortions. These resampled BOLD time-series will be referred to as preprocessed BOLD in original space, or just preprocessed BOLD. The BOLD time-series were resampled into standard space, generating a preprocessed BOLD run in [‘MNI152NLin2009cAsym’] space. First, a reference volume and its skull-stripped version were generated using a custom methodology of fMRIPrep. Several confounding time-series were calculated based on the preprocessed BOLD: framewise displacement (FD), DVARS and three region-wise global signals. FD and DVARS are calculated for each functional run, both using their implementations in Nipype (following the definitions by Power et al. 2014). The three global signals are extracted within the CSF, the WM, and the whole-brain masks. Additionally, a set of physiological regressors were extracted to allow for component-based noise correction (CompCor, Behzadi et al. 2007). Principal components are estimated after high-pass filtering the preprocessed BOLD time-series (using a discrete cosine filter with 128s cut-off) for the two CompCor variants: temporal (tCompCor) and anatomical (aCompCor). tCompCor components are then calculated from the top 5% variable voxels within a mask covering the subcortical regions. This subcortical mask is obtained by heavily eroding the brain mask, which ensures it does not include cortical GM regions. For aCompCor, components are calculated within the intersection of the aforementioned mask and the union of CSF and WM masks calculated in T1w space, after their projection to the native space of each functional run (using the inverse BOLD-to-T1w transformation). Components are also calculated separately within the WM and CSF masks. For each CompCor decomposition, the k components with the largest singular values are retained, such that the retained components' time series are sufficient to explain 50 percent of variance across the nuisance mask (CSF, WM, combined, or temporal). The remaining components are dropped from consideration. The head-motion estimates calculated in the correction step were also placed within the corresponding confounds file. The confound time series derived from head motion estimates and global signals were expanded with the inclusion of temporal derivatives and quadratic terms for each (Satterthwaite et al. 2013). Frames that exceeded a threshold of 0.5 mm FD or 1.5 standardized DVARS were annotated as motion outliers. All resampling can be performed with a single interpolation step by composing all the pertinent transformations (i.e. head-motion transform matrices, susceptibility distortion correction when available, and co-registrations to anatomical and output spaces). Gridded (volumetric) resampling were performed using antsApplyTransforms (ANTs), configured with Lanczos interpolation to minimize the smoothing effects of other kernels (Lanczos 1964). Non-gridded (surface) resamplings were performed using mri_vol2surf (FreeSurfer). Many internal operations of fMRIPrep use Nilearn 0.5.2 (Abraham et al. 2014, RRID:SCR_001362), mostly within the functional processing workflow. For more details of the pipeline, see the section corresponding to workflows in fMRIPrep's documentation.
